# Forced swim stress dynamically alters neural activity and bi-directionally modulates NMDA receptors in the prefrontal cortex

**DOI:** 10.1101/2021.03.10.434848

**Authors:** Marc-Alexander L.T. Parent, Amber Lockridge, Li-Lian Yuan

**Author notes:** Address correspondence to Dr. Li-Lian Yuan, Department of Physiology and Pharmacology, Des Moines University, Des Moines, IA 50132, USA.

## Abstract

Repeated exposure to stress results in progressively divergent effects on cognitive behaviors that are dependent on the integrity of networks in the medial prefrontal cortex (mPFC). To investigate molecular mechanisms responsive to variable repetition of mild stress, we measured persistent neural activity, *in vitro*, from mPFC slices in mice that had been repetitively exposed to 10 minutes of forced swim stress for 3-10 days. 3-day short-term stress facilitated persistent neural activity by increasing event duration while 10 days suppressed event duration and amplitude. These dynamic changes were accompanied by a similar bi-directional modulation of the NMDA/AMPA receptor current ratio, an important synaptic mechanism for sustaining the persistency of neural activity. Specifically, short-term stress led to potentiated NMDA currents with slower decay kinetics, and extended stress produced smaller currents with faster decay. The inhibitory action of ifenprodil, a specific blocker of NR2B-containing NMDA receptors, was more effective in NMDA current suppression following light stress and less effective after longer stress compared to naive controls. Persistent activity and glutamate receptor balance in the neocortex have been linked to working memory and impulse control. Therefore, these results could provide insight for generating therapeutic strategies to prevent or reverse stress-induced cognitive deficits.

## Introduction

One dramatic effect of an initial encounter with a stressful stimulus is the rapid shunting of cognitive processing toward a more reflexive and/or instinctive fight-or-flight response. This is a widely accepted psychological principle, but it may overshadow many of the finer nuances of the body’s stress response over time and the impact of that response on behavior. Performance of cognitive tasks is negatively affected by concurrent stress or stress occurring shortly before (Arnsten and Goldman-Rakic 1998; Holmes and Wellman 2009; Liston, McEwen et al. 2009). Repetitive exposure to stress also leads to structural changes including neuronal spine loss and dendritic atrophy(Cook and Wellman 2004; Radley 2005; Goldwater, Pavlides et al. 2009). However, it has also been reported that acute, moderate stress facilitates classical conditioning and associative learning (Shors, Weiss et al. 1992; Joëls, Pu et al. 2006b) as well as improving performance on a working-memory task (Yuen, Liu et al. 2009). These converse data highlight the dynamic nature of the brain’s response to stress as well as the need for development of hypotheses questioning how these divergent effects are implemented at the level of local prefrontal cortical (PFC) circuits.

Pyramidal neurons located in the PFC receive tens of thousands of synaptic inputs and can undergo a transition membrane potential transition from hyperpolarized to a prolonged depolarization state. This type of neuronal persistent activity, which may last for several seconds, represents a synchronous activation of sub-networks in the PFC region and has been proposed as a candidate cellular mechanism for how an organism holds information within a neural network to guide behaviors (Goldman-Rakic 1995; Fuster 1998; Funahashi 2001; Miller and Cohen 2001). Previous studies have obtained recordings of persistent activity from *in vitro* slices but at a characteristically low spontaneous frequency (Sanchez-Vives, McCormick et al. 2000; Tseng 2004). Various methods have been employed to increase the probability of obtaining these events, including organotypic culture (Tu, Kroener et al. 2007), acute slices from an early developmental stage (Cossart, Aronov et al. 2003), application of dopamine (Tseng 2004), specific species preparations such as ferret (Shu, Hasenstaub et al. 2003) and by modifying ionic composition of bath solution (Steriade, Contreras et al. 1993).

In this investigation, we recorded spontaneous bouts of persistent activity using standard aCSF over long period of time to characterize events associated only with the strongest local recurrent networks retained in the slice, without the influence of bath applied neuromodulators or altered ionic contents of recording solution. Spontaneous persistent activity recorded in this way shows similarity to those identified *in vivo*, based on duration, kinetics of onset and decay (Seamans 2003; Haider 2006; Gruber and O’Donnell 2009) as well as to PA events reported in previous studies done in normal aCSF *in vitro* (Tseng 2004). We used this technique combined with various amount of forced swim exposure to mice to investigate whether varying durations of repetitive stress differentially modulate mPFC network activity and through what molecular mechanisms these effects are achieved.

## Materials and Methods

### Animals and Behavioral Stress

10-to 15-week-old male mice (129 SvEv) were used in this study. For mild stress exposure, mice were placed in clear plexiglass cylinders (20 cm tall x 15 cm wide) filled with tap water to a height of 14 cm and a temperature of 25±1°C for 10 minutes followed by 10 minutes in a dry warming cage (∼27°C). This was repeated for 3 or 10 consecutive days with control mice for each treatment group handled briefly for a similar number of days. 24 hours following the last exposure, mice were sacrificed for *in vitro* recording. The use of animals for studies described here was approved by the University of Minnesota Institutional Animal Care and Use Committee.

### PFC Slice Preparation

The experimenter, performing whole-cell recordings, was kept blind to pretreatment condition during both data acquisition and data analysis. After exposure to behavioral stress, mice were anesthetized by a lethal dose of mix of ketamine/xylazine and perfused through the heart with ice-cold cutting solution containing (in mM) 240 Sucrose, 2.5 KCl, 1.25 NaH2PO4, 25 NaHCO3, 0.5 CaCl2, 7 MgCl2, and 7 Dextrose. 300 μm coronal slices of PFC were prepared according to Parent et al. (Parent, Wang et al. 2009).

### Electrophysiology

After incubation in normal aCSF (in mM): 125 NaCl, 2.5 KCl, 1.25 NaH2PO4, 25 NaHCO3, 2 CaCl2, 1 MgCl2, 10 Dextrose at room temperature, slices were transferred into the recording chamber under a Zeiss Axioskop 2, fitted with 40X water-immersion objective and differential interference contrast (DIC). Infrared light, in conjunction with a contrast-enhancing camera, was used to visualize individual pyramidal neurons in the prelimbic (PrL) region of PFC. Data were acquired through Axograph X software and a Multiclamp 700B amplifier. For spontaneous persistent activity recording, aCSF was slightly modified, according to Cossart et al (2003), containing (in mM) 123 NaCl, 3 KCl, 26 NaHCO3, 1 NaH2PO4, 2 CaCl2, 2 MgSO4 and 10 dextrose. Pipette solution contained (in mM) 120 K-gluconate, 20 KCl, 10 HEPES, 0.2 EGTA, 2 MgCl2, 4 Na2-ATP, 0.3 Tris-GTP, and 14 Phosphocreatine. For voltage clamp recordings of evoked EPSCs, a mono-polar stimulation electrode was placed ventral and perisomatically (∼75 to 100 □m away) from patched neurons within the same deep layers of the prelimbic region. 100 μM picrotoxin was added to the aCSF to block GABAa receptor mediated IPSCs. Pipette solution contained (in mM) 117 cesium gluconate, 2.8 NaCl, 20 HEPES, 0.4 EGTA, 5 tetraethylammonium-Cl, 2 MgATP, and 0.3 MgGTP, and 8 QX314, pH 7.2–7.4 (285–295 mOsm). In determining relative NMDAR and AMPAR contributions to total glutamate current, 50 μM D,L-APV was added to the aCSF to block the NMDA component. To evaluate NR2B-mediated NMDA currents, 3 μM ifenprodil was added in the presence of 20 μM CNQX and 100 μM picrotoxin. To record miniature EPSCs, 500 nM TTX was added to aCSF to block Na^+^ channels.

### Data Analysis and Statistics

Because of the large amount of time-series data, we built a set of scripts in MATLAB for automated detection and analysis of neural persistent activity in order to add objectivity and reduce labor (Parent and Yuan 2011). Detected events were subjected to subsequent visual confirmation by the experimenter. Three neurons exhibiting high frequency firing were excluded because detection of discrete events in these neurons was more difficult and increased the probability of incorporating false positives into the data set. NMDA current curve fitting and quantification of evoked current recordings were done using the analysis routines contained in Axograph X. Miniature EPSCs were analyzed in MiniAnalysis (Synaptosoft). All graphing and statistics were carried out in OriginPro 8.5 (Originlab Corporation). Significance (p < 0.05) was determined by one-way ANOVA. Error bars represent SEM.

### Drugs

APV, CNQX, picrotoxin, QX314, and ifenprodil were purchased from Tocris, and TTX from Alomone.

## Results

To investigate changes in neural activity in response to varied stress duration, we recorded events from layer 5 pyramidal neurons in the dorsal prelimbic (PrL) cortex (Fig. 1AB) since these cells have been shown to receive local recurrent synaptic inputs, show persistent activity in vivo (Lewis and O’Donnell 2000; Seamans 2003; Haider 2006), and are vulnerable to chronic stress (Goldwater, Pavlides et al. 2009). These pyramidal neurons were identified using the following criteria: proximity to the dorsomedial bend of the cingulum/corpus callosum as it moves laterally over the striatum at approximately 1.7mm anterior to bregma, pyramidal shaped soma and major apical dendrites, and the regular spiking pattern with slight adaptation in response to sustained current injections (Fig. 1AB). Extended 30 minute recording durations were used in the absence of extracellular stimulation and receptor blockers to compensate for low *in vitro* spontaneous persistent activity. According to our standards, membrane potential depolarizations were grouped as persistent activity when the depolarizations were larger than 2 mV, lasting for more than 500 ms, and exhibiting slower onset and decay than that of excitatory post-synaptic potentials (EPSPs). The majority (∼88%) of recorded events across all group conditions occurred without reaching the firing threshold, while 12% did drive neurons to fire action potentials (Fig. 1C). Using this preparation, we were able to demonstrate modulation of persistent activity *in vitro* by manipulating network excitability and synaptic connectivity. For example, decreasing aCSF magnesium concentration to 0.1mM dramatically increased the occurrence of spontaneous persistent events (Fig. 1D) without depolarizing the membrane potential of neurons, suggesting low Mg^2+^ may exert influences through synaptic mechanisms.

**Figure 1.**
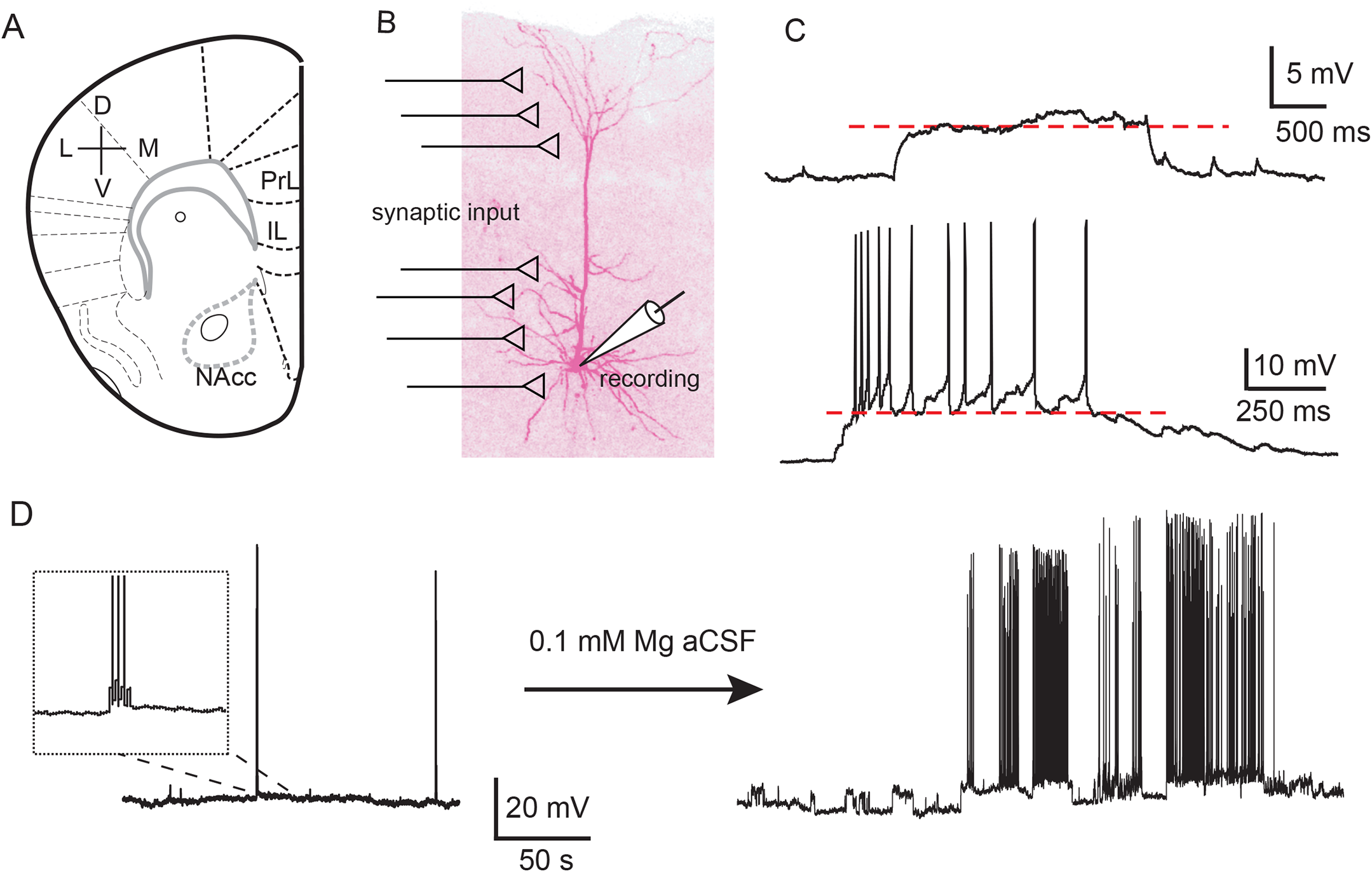
Recording persistent neural activity in mPFC slices. (**A**) Coronal slice preparation containing the medial PFC (PrL: prelimbic and IL: infralimbic regions). All recordings were made in PrL. (**B**) A layer 5 pyramidal neuron filled with biotin. (**C**) Spontaneous persistent activity with (upper) or without (lower) concurrent firing, recorded in the absence of receptor blockers. (**D**) Lowering Mg^2+^ concentration to 0.1 mM in aCSF, a manipulation that promotes NMDAR functioning dramatically increased the level of persistent activity.

### Repetitive stress exposure induced dynamic changes in persistent activity

We gave mice up to 10 consecutive days of 10-minute forced swim (FS) (Porsolt, Le Pichon et al. 1977). Tissue from 2 treatment groups (3 and 10 days) was dissected 24 hours after the last stress to make coronal slices of the mPFC for recording (Fig. 2A). Treatment time points were correlated to major increases in immobility behavior, a potential reflection of the subject’s shift in perception and response strategy to the stress condition (Thierry, Steru et al. 1984; Lockridge, Su et al. 2010).

**Figure 2.**
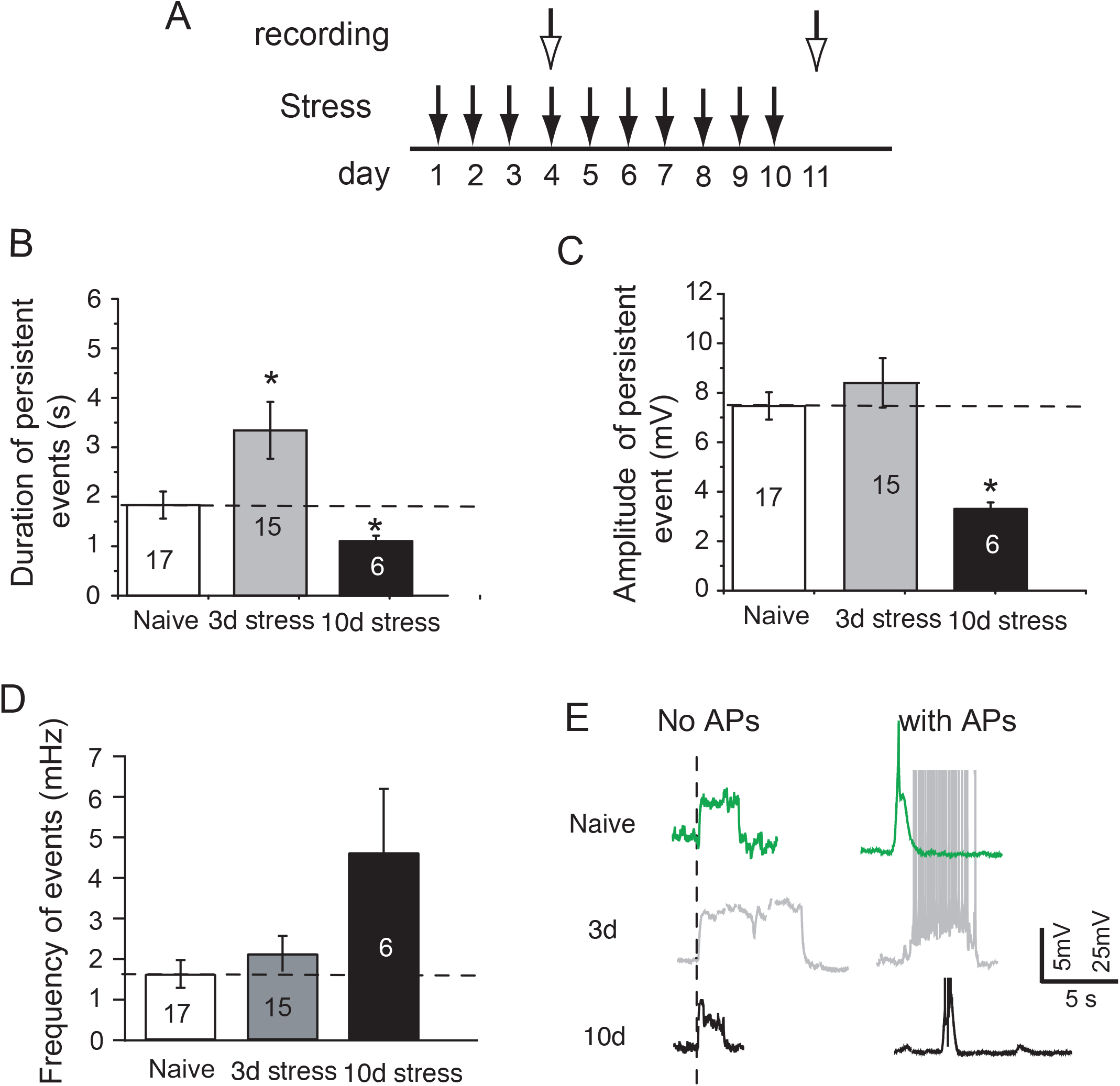
Dynamic changes in persistent activity of mPFC pyramidal neurons induced by progressive increase in the number of exposures to swim stress. (**A**) Experimental design of behavioral stress. 10 minutes forced swim (FS) stress was administered up to 10 consecutive days with groups removed for recording after 3 days (3d) or 10 days (10d). (**B**) Representative traces of persistent activity with (right) and without (left) action potential (AP) firing under various conditions. (**C**) Summary of duration and amplitude of persistent activity recorded under various stress conditions: naïve (n=17 cells), 3-days (n=15) and 10-days stress (n=6). *p<0.05 vs. naïve control.

Compared to naïve mice, increasing stress exposure induced a biphasic change in persistent activity level (Fig. 2; n=17 cells/N=13 animals, 15/N=10, and 6/N=3 recorded for naïve, 3 days, and 10 days stress). The mean duration of events recorded from naive mice was 1.83±0.27 seconds. Exposure to 3-day stress significantly increased the duration to 3.34±0.57 sec (p<0.05). By contrast, after 10 days of stress, persistent event duration was diminished by 40% to 1.09±0.14 sec (p<0.05) (Fig. 2B). Event amplitude in this group was also suppressed from 7.47±0.55 mV (naïve) to 3.26±0.24 mV (p<0.05) but remained largely unchanged in 3-day stressed animals (8.4±0.39 mV; Fig. 2C). Frequency of persistent activity events was not significantly different across all groups (naive: 1.63±0.34 mHz; 3d: 2.13±0.44 mHz; 10d: 4.63±1.57 mHz).

### Are intrinsic ionic mechanisms targeted by stress?

We first investigated whether stress-induced alterations in intrinsic membrane properties could contribute to the dynamic changes in persistent activity duration we observed. The resting membrane potential was close to -70 mV, relatively hyperpolarized compared to typical *in vivo* conditions (Yang, Seamans et al. 1996; Dégenètais, Thierry et al. 2002) but showed no difference across groups (Naive: -69.00±.0017 mV, n=17; 3d-FS: -65.4±.0018 mV, n=15; 10d-FS: -68.48±.0018 mV, n=6). Hyperpolarization-activated HCN channels could theoretically exert influence on both membrane excitability and synaptic integration. However, we measured HCN-mediated voltage sag in response to a negative current injection (−150 pA) and found no significant difference in the sag amplitude among variously stressed groups (naïve: 1.5±0.1mV, n=50 cells/N=17 mice; 3d: 1.8±0.2 mV, n=43/N=18; 10d: 1.7±0.3 mV, n=10/N=5) (Fig. 3B). Input resistance following current injection or rheobase showed no significant difference between groups (Fig. 3CD). Changes in the expression of after-hyperpolarization potentials (AHP) could also regulate the duration of persistent activity. A burst of 10 action potentials yielded post-burst AHPs consisting of two primary components: an initial medium (mAHP) occurring within the first 500 ms of the final action potential of the burst followed by a prolonged slow AHP (sAHP) that extended for hundreds of milliseconds after the mAHP (Fig. 3E). Interestingly, we found that the mAHP amplitude of neurons in the 10d stress group was significantly larger (−2.40±1.16 mV, n=10/N=5, p<0.05) than in the naïve (−1.58±.82 mV, n=38/N=17) or 3-day stress groups (−1.66±1.25 mV, n=26/N=15) (Fig. 3F). sAHP, as measured by the area under the curve starting 500 ms after the final action-potential, remained similar across all groups (Fig. 3G).

**Figure 3.**
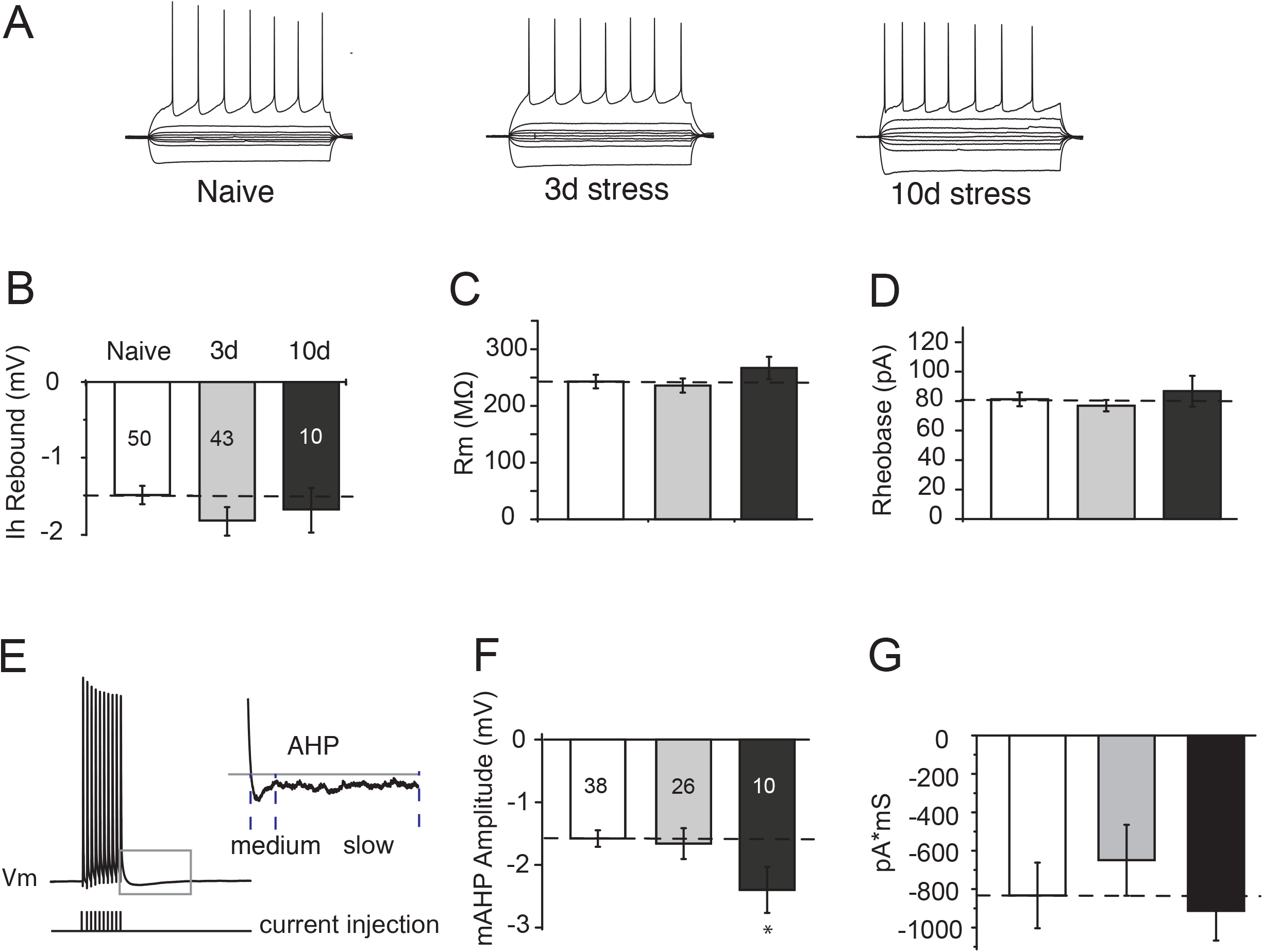
Effects of repetitive stress on intrinsic membrane properties of mPFC pyramidal neurons. (**A**) Representative traces of neural responses to a family of depolarizing and hyperpolarizing current injections. (**B**) Amplitude of Ih-based membrane voltage (Vm) rebound as an indicator of HCN channel involvement in current form modulation showed no difference between groups (n=50, 43, and 10 for naïve, 3d-, and 10d-stress groups). Input resistance (Rm) (**C**) and rheobase (**D**) showed no difference among conditions (naïve: n=50/52, 3d: n=43/41, 10d: n=10/10 cells). (**E**) An example of after hyperpolarization potential (AHP) components, following a current injection triggered burst of 10 action potentials, consisting of an initial medium phase (mAHP) followed by a slow phase (sAHP). (**F**) mAHP in the 10d-FS group (n=10), calculated from the initial 500ms of an activity-driven AHP following 10 current injections, was increased relative to the naïve group (n=38) and 3d-FS (n=26). *p<0.05. (G) sAHP, measured by area under the latter portion of the curve, was not significantly impacted by treatment.

### Biphasic changes of glutamatergic transmission

We next investigated synaptic mechanisms as an alternative to explain stress-related biphasic changes in persistent activity. In the presence of picrotoxin (100 μM) to block inhibitory currents, we first recorded total glutamate currents (EPSC) in response to electrical stimulation from neurons voltage-clamped at -70 mV. After establishing stable monosynaptic responses, the holding voltage was brought to +40 mV to augment the NMDA-mediated portion of the evoked glutamate current. Once a new stable baseline EPSC containing both AMPA and NMDA contributions was achieved, 50 mM D,L-APV was added to the aCSF to block the NMDA component of the total glutamate current. The remaining post D,L-APV application EPSC contained only the AMPA-mediated portion of the total glutamate current. We computed the NMDA-mediated portion of the total glutamate current by subtracting the average AMPA component from the average total evoked glutamate current (Fig. 4A). Both peak current amplitude and the total charge are indicators of synaptic efficacy of pyramidal neurons in mPFC. Since NMDA current size is dependent on stimulation intensity, we normalized it to the AMPA currents evoked from the same synapses (Ngo-Anh, Bloodgood et al. 2005) and used changes in the NMDA/AMPA current ratio, either in peak amplitude or current area, as indicators of synaptic plasticity (Thomas, Beurrier et al. 2001; Myme 2003).

**Figure 4.**
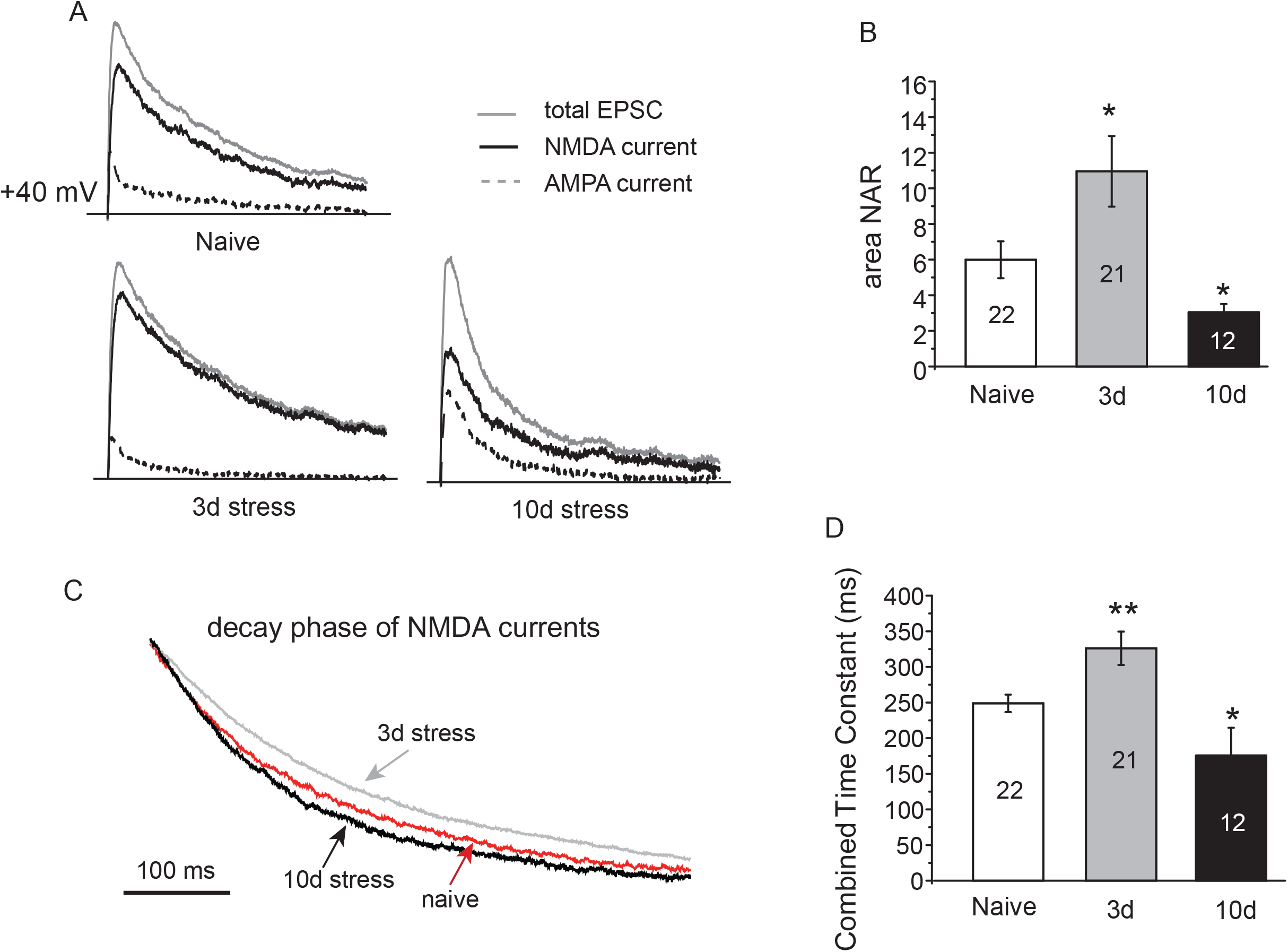
Biphasic changes of NMDA/AMPA current ratio. (A) Separation of NMDA and AMPA currents from total EPSC by pharmacological tools. (B) Trace averages showing the decay phase of NMDA currents after exposure to the different stress conditions. (C) NMDA/AMPA current ratio was quantified by total charge, i.e. integral area underneath the current, and compared among various groups. (D) Statistical comparison of NMDA current decay time constant (τ). N=22, 21,12 for naïve, 3d-, and 10d-stress groups. *p<0.05; **p<0.01 vs. naïve control.

While a trend of increase in peak *amplitude* NAR in 3d-group was not statistically significant (naïve: 2.4±0.2, n=22/N=13; 3d: 3.0±0.4, n=21/N=13), the NAR of 10d-group was reduced to 1.5±0.2, (n=12/N=6, p<0.05). A close inspection of individual current traces (Fig. 4A) indicated NMDA current decay kinetics may be involved as a factor that could influence total charge and not amplitude (Fig. 4A). *Area-*based NAR changes subsequently revealed a significant 80% increase in 3d- and a 45% decrease in 10d-FS groups relative to controls (naïve: 6.0±1.0; 3d: 11.0±2.0; 10d-FS: 3.0±0.5) (Fig. 4B). These results suggest that NMDA receptors contribute significantly to evoked excitatory currents and highlight the impact of NMDA current decay on glutamate neurotransmission efficacy.

We further quantified current decay by calculating a weighted decay time constant (τ) for isolated NMDAR current using the formula τ = ((A1*T1)+(A2*T2)) and plugging in the computed least-square values from a fit double exponential decay equation. Again, stress with different duration exhibited differential effects. Current decay was 31% slower (326.2±23.4 ms, p<0.01) and 29% faster (175.6±39.0 ms, p<0.05) in neurons from the 3d-FS and 10d-FS groups, respectively (Fig. 4D). These stress-responsive biphasic changes in decay kinetics pointed towards NMDA receptor (NMDAR) composition shifts.

### NR2B subunit contribution to NMDA currents was targeted by stress

NR2B-containing NMDARs are known to exhibit slower current decay properties compared to NR2A-containing receptors (Monyer, Burnashev et al. 1994) prompting us to investigate stress-induced NMDAR subunit composition changes. We isolated NMDA currents with CNQX (AMPA blocker; 20 mM) and picrotoxin (100 mM) and calculated percentage current inhibition after application of the NR2B antagonist ifenprodil (3 μM) (Fig. 5A). The 3d-group showed a significant increase in NR2B-related current inhibition by ifenprodil with enhanced suppression of current amplitude from 46.8±6.0% (naïve, n=9/N=3) to 62.7±3.1% (3d, n=8/N=2) and area under the curve from 53.5±5.7% naïve to 72.9±3.9% 3d (p<0.05). Conversely, 10d-group currents were less inhibited by ifenprodil (amplitude: 26.8±4.8%; area: 30.1±6.2%; n=6/N=3; p<0.05 vs. naïve) (Fig. 5B). Taken together, the proportion of functional NR2B subunit containing NMDARs at the synapse appeared to undergo similar dynamic changes in response to varying repetitive stress as the NMDA current kinetics, paralleling functional changes in persistent event duration.

**Figure 5.**
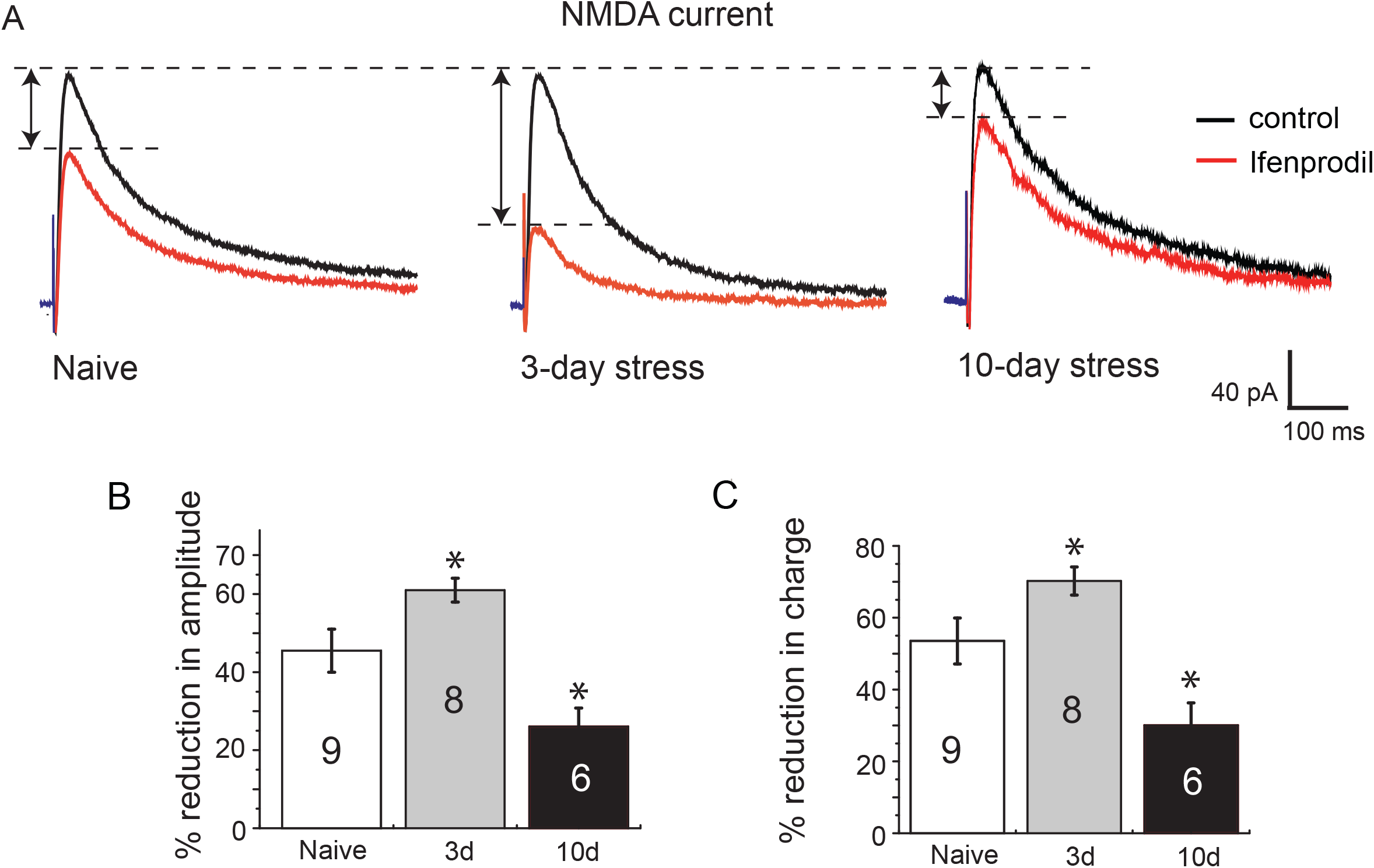
Contribution of NR2B-containing receptors to total NMDA currents. (A) Isolation of NR-2B component using a NR-2B specific blocker ifenprodil by subtracting ifenprodil (grey) from total NMDA currents (black). (B) Quantification of percentage reduction in current amplitude by ifenprodil. (C) Quantification of ifenprodil –induced percentage reduction in total charge carried by NMDA currents. N=9, 8, and 6 for naïve, 3d- and 10d-stress groups. *p<0.05 vs. naïve control.

### AMPA receptor-mediated miniature EPSCs

While the measured decay kinetics highlight changes associated with NMDA receptors, the question of whether the AMPA component, as a factor in the NMDA/AMPA current ratio, is static at different phases of stress became pertinent. Previous studies showed that a single exposure to acute swim stress increased AMPAR-mediated miniature EPSC (mEPSC) amplitude without affecting presynaptic glutamate release (Yuen et al., 2009). We examined mEPSCs at two further time points: 3d and 10d, in the presence of 500 nM TTX to block action potential evoked synaptic transmission (See Fig. 6). There was no obvious difference in the mean mEPSC amplitude between groups (naïve: 9.8±1.0 pA, n=9/N=7; 3d: 9.9±0.8 pA, n=8/N=6; 10d: 10.9±0.4 pA, n=6/N=2) (Fig. 6B). However, 3d-FS and 10d-FS groups showed an 80% increase and a 70% decrease, respectively, in mEPSC event frequency (naïve: 1.04±0.26 Hz; 3d: 1.87±0.23 Hz; 10d: 0.33±0.08 Hz, p<0.05) (Fig. 6C). These data suggest that repetitive stress does not alter AMPAR-mediated postsynaptic responses, but presynaptic glutamate release at the 3d and 10d stress regimes underwent opposite changes. The direction of these changes is consistent with that of persistent activity following similar times of stress exposure.

**Figure 6.**
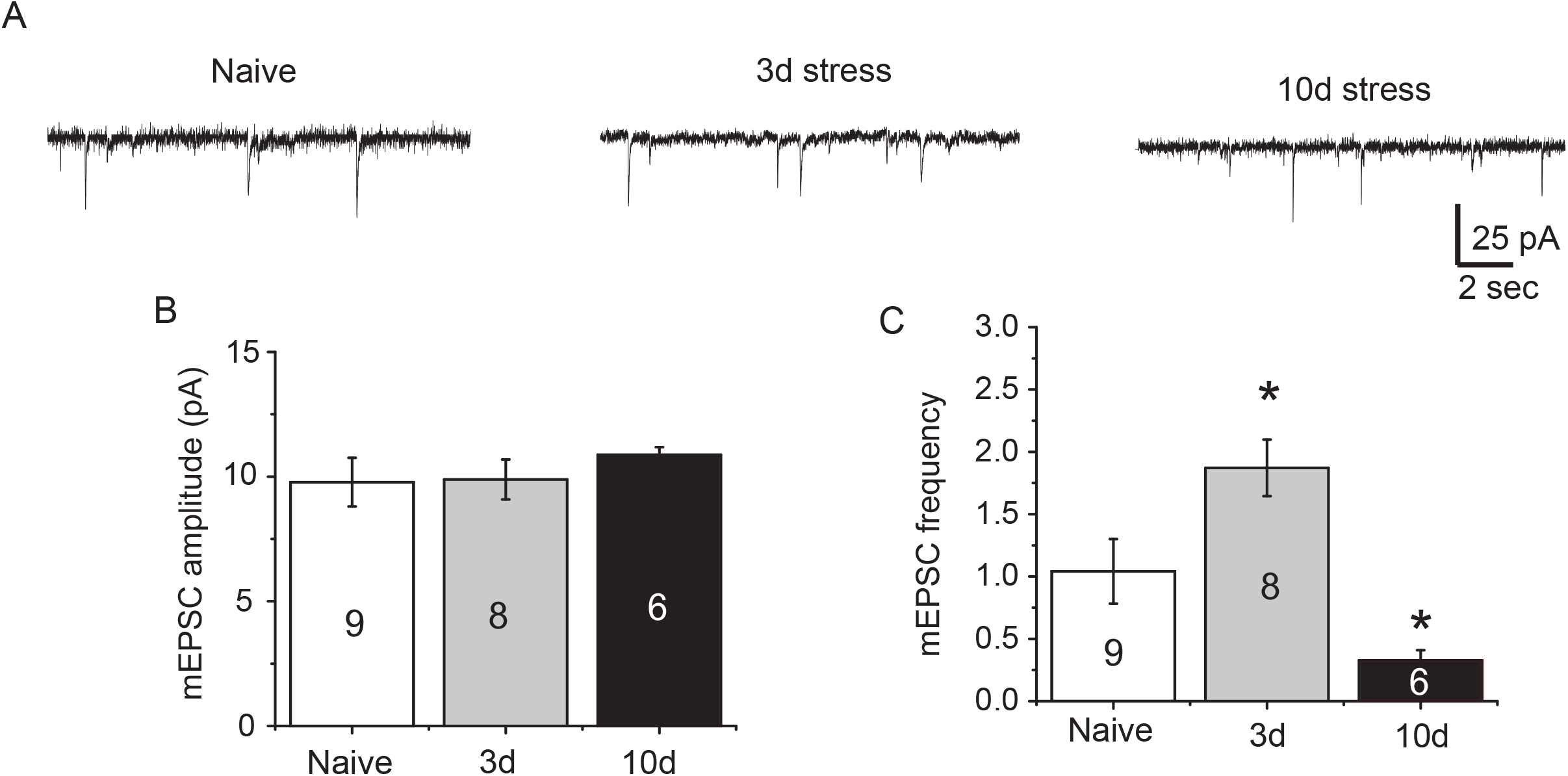
Miniature EPSC analysis of AMPA receptor-mediated synaptic transmission. (A) Sample traces showing mEPSCs (downward deflections) recorded at -70 mV from each group: naïve (n=9), 3d (n=8), and 10 day stress (n=6). (B) Stress did not show a significant effect on mEPSC amplitude. (C) mEPSC frequency was altered, in opposite directions, by short-term and chronic stress exposure. * p<0.05 vs. naïve control.

## Discussion

### Biphasic effects of repetitive stress on neural activity

Network-driven persistent neural activity in the PFC has been proposed as a potential neurological correlate of working memory processes (Batuev, Kursina et al. 1990; Goldman-Rakic 1996; Fuster 1998; Baeg, Kim et al. 2003). Its occurence in the neocortex is thought to originate from a re-entrant collateral network of local excitatory pyramidal neurons and has been observed both *in vivo* (Steriade, Contreras et al. 1993; Branchereau, Van Bockstaele et al. 1996; Lewis and O’Donnell 2000; Seamans 2003; Haider 2006) and *in vitro* (Sanchez-Vives, McCormick et al. 2000; Cossart, Aronov et al. 2003; McCormick 2003; Tseng 2004) in the PFC. In this study, we monitored the effects of repeated stress on spontaneous persistent activity *in vitro*, which revealed a biphasic modulation taking place at molecular, cellular, and network level. Mild acute and repeated stress enhanced neocortical network activity in mice but repetition beyond a critical threshold reversed direction and drove persistent activity down. Concomitantly, short- and long-term stress respectively increased and decreased the NMDA/AMPA ratio property of excitatory synapses. This can be attributed, at least in part, to bidirectional shifts in NMDAR function and/or receptor subunit composition. This hypothesis was supported by the involvement of NR2B-mediated subunit composition changes.

While biphasic modulation by varying repetitions of stress exposure has been reported, these studies have focused on neural structures. For example, acute swim stress increased hippocampal benzodiazepine receptors in rats while prolonged repeated exposure reduced the numbers in the same area (Avital, Richter-Levin et al. 2001). The second study used daily injections of corticosterone, a central biomarker of stress exposure. Short-term corticosterone increased the concentration of hippocampal neural cytoskeletal proteins while long-term administration diminished protein levels (Zhao, Xie et al. 2009). The location of these changes in the hippocampus is particularly interesting since PFC persistent activity is dependent on intact hippocampal afferents, though this input cannot independently drive persistent activity (Lewis and O’Donnell 2000).

### Stress-induced dynamic changes in glutamatergic transmission

Both ionic and synaptic mechanisms have been implicated in the generation and maintenance of persistent activity (Major and Tank 2004). NMDA receptors, owing to their voltage and ligand dependence, as well as slow kinetics, represent a synaptic mechanism for the fine-tuning of persistent activity. Computation modeling (Wang 1999; Durstewitz, Seamans et al. 2000), *in vivo* (Steriade, Contreras et al. 1993) and *in vitro* studies (Schiller, Major et al. 2000; Milojkovic 2005) have demonstrated that the synaptic drive needed to maintain persistent activity is likely derived from NMDA currents (Antic, Zhou et al. 2010). Furthermore, the synaptic NMDA/AMPA current ratio (NAR) in the mPFC region may play a significant role in translating the immediate biological consequences of stress into changes in glutamatergic transmission efficacy that mediate persistent activity. Recent studies discovered that single, acute exposure to mild stress enhances working memory and glutamatergic transmission, possibly through serum- and glucocorticoid-inducible kinase (SGK) dependent mechanisms that facilitate AMPA and NMDA receptor trafficking (Yuen, Liu et al. 2009; Yuen, Liu et al. 2010).

In the case of short-term and chronic stress, our results suggest NAR, and NMDA receptors in particular, do serve as an effector of stress to influence PFC activity and function. The 10d-FS showed significant decreases in NAR indicating reduced NMDAR participation in driving synaptic current efficacy. Elevated NAR values for the 3d-FS group were most significant when calculated by area rather than amplitude, implicating the likely involvement of current decay kinetics. In fact, the decay time constant was enhanced after 10d stress and diminished by less stress exposure, moderately in the 3d condition. A prolonged decay phase in the moderately stressed mice accounts for stress-induced increases in area-based NARs. A slower decay of NMDA currents during glutamatergic neurotransmission inevitably indicates an increased entry of charge into neurons during excitatory synaptic activity. This may lengthen post-synaptic depolarizations of neurons following presynaptic glutamate release. On the other hand, the apparent absence of changes to NAR in the 1d condition is at odds with the finding that this group showed the largest increase in persistent activity duration. How can this be reconciled? In the current study, both NMDA and AMPA receptor-mediated currents were evoked by electrical stimulation, representing activity from the same set of synapses on a given neuron. Therefore, the NMDA/AMPA current ratio holds to a population of synapses. Logically, an increase in both components would yield slight or no change in their ratio. Previous studies have shown that a single exposure to acute swim stress increases AMPAR-mediated miniature EPSC (mEPSC) amplitude without affecting presynaptic glutamate release (Yuen, Liu et al. 2009). Our own testing indicated no changes in AMPAR-mediated postsynaptic responses for 3d and 10d stressed animals. It is likely, therefore, that NMDAR enhancement after 1 day of stress was still a primary contributor to elevated persistent activity durations but was balanced in the NAR calculation by similar AMPAR increases only within the 24-hour time frame.

### Stress-induced changes in NMDAR subunit composition

NMDARs in the neocortex are predominantly composed of NR1/NR2A and NR1/NR2B subtypes, and these subtypes have distinct decay rates. NMDA currents at the recurrent synapses of PFC in adult rats exhibit a longer decay time constant (τ) than other cortical regions, resulting in more effective temporal summation (Wang, Stradtman et al. 2008). This can be attributed to enrichment of NR2B subunits in these synapses as NR2B-containing NMDAR mediated currents are relatively slow in decay kinetics (Monyer, Burnashev et al. 1994). Using the NR2B-specific antagonist ifenprodil, we found a larger fraction of NR2B-containing NMDARs contributing to evoked glutamatergic neurotransmission in 3d-FS group. This finding supports the notion that moderate repetition to stress exposure enhances the expression of receptors with this subunit. Lack of change in current decay rate by 1-day stress indicates that a subunit composition shift may be achieved via slow processes such as gene expression or relatively long-distance translocation of receptors. On the other hand, 10d-FS tissue was less responsive to ifenprodil, indicating a reduction in NR2B-containing NMDARs and suggesting that changes in subunit expression or trafficking are more dynamically responsive to stress. It is unclear where the threshold for reversal occurs but we speculate between 5-7 days in mice.

### Other mechanisms involved in stress effects on persistent activity

Clearly, a large number of signaling pathways and molecules are targeted by repetitive exposure to stress (Robbins and Arnsten 2009). We thus cannot rule out that other effectors besides NMDA receptors contribute to the biphasic changes in persistent activity we observe. Inactivation of HCN channels in PFC prolongs persistent activity *in vitro* and strengthens functional connectivity of networks *in vivo* (Wang, Ramos et al. 2007), whereas activation of SK channel-mediated after-hyperpolarization potentials (AHPs) terminates persistent activity and shunts excitatory neurotransmission (Hagenston, Fitzpatrick et al. 2007; Faber 2010). We found that stress did impact several neural intrinsic properties but not in a biphasic manner within the time frames of treatment we used. Changes to input resistance and rheobase predicted increased excitability for the 1d-FST group but repetitive stressing normalized these to naïve values. The initial mAHP for 10d-FS treated neurons, occurring within the first 500 ms after the action potential train, was greater in amplitude than in other groups, which could participate in the shunted duration of persistent activity. It is particularly worth noting, however, that HCN channels, a key component connecting stress, neural activity, and working memory (Wang, Ramos et al. 2007), did not appear to be changed by stress in our experiments. It is possible that the neuromodulatory tone that influences neural circuits *in vivo* through HCN channels is not maintained in slice.

### Molecular Model of cortical network activity under stress

Together, our results suggest that dynamic changes in glutamatergic transmission induced by repetitive stress can be depicted by the following model. Both AMPA and NMDA currents are up-regulated following acute or short-term exposure to stress. The trending increase in NMDA/AMPA ratio indicates initial exposure to stress may enhance the NMDA component slightly more than the AMPA component; however both components are augmented at this early phase. While NMDAR-mediated transmission remains facilitated, AMPAR-mediated transmission quickly returns to baseline while presynaptic release probability remains high. Continued daily exposure to stress for an extended period of time ultimately reverses the enhancement of NMDA receptors in the mPFC, yielding a dramatic decrease in the NMDA contribution to excitatory neurotransmission, a decrease in presynaptic release probability, and an increase in activity-driven AHPs. Our data suggest that the changes in the NMDA component to neurotransmission in mPFC are largely mediated by subunit switching, specifically an initial increase in functional NR2B subunit-expressing NMDA receptors at mPFC synapses followed by a decrease. It remains to be answered, however, whether mechanisms involved in acute stress sustain or accumulate through the repetitive exposure or whether additional players are recruited during the subsequent exposures and new mechanisms are required to drive the reversal phenomenon.

## Acknowledgements

We would like to thank Drs. William Engeland and Matthew Chafee for insightful discussions on the project and members of the Yuan lab for help with the behavioral tests including Arman Cicic, Anja Srienc and Cassie Kowalski.

